# Macrophage metabolism directs regenerative versus fibrotic healing through BMP signaling in the mouse digit tip

**DOI:** 10.64898/2026.05.04.722661

**Authors:** Mimi C. Sammarco, Siqi Liu, Ni Su, Madhumidha Ramesh, Charlotte Raymond, John Carleton, Anne Le, Alexander J. Trostle, Robert J. Tower, Jennifer Simkin

## Abstract

Macrophages play a central role in determining the outcomes of healing, coordinating regeneration in some injuries and scar formation in others. In both cases, this coordination involves the cross-talk between macrophages and surrounding cells. But what drives the different cross-communication pathways to determine healing outcomes is not well known. In this study, we make use of the mouse digit tip amputation model, in which an amputation through the third phalangeal element (P3) is able to completely regenerate whereas an amputation through the second phalangeal element (P2) forms a scar. We identify a population of macrophages that is specific to the P3 regenerating digit. By integrating single-cell RNAseq, spatial transcriptomics, and metabolomic analyses, we show that this population localizes specifically to the growing bone front, express BMP ligands that drive downstream BMP activation in neighboring osteoblasts and is governed by a two-part metabolic switch involving increased fatty acid oxidation coupled with reduced glycolytic activity. This spatially restricted, BMP-expressing macrophage population is entirely absent in the scar-forming P2 injury, and our data indicate that environmental conditions unique to the regenerating digit are responsible for its emergence. Together these findings identify a regeneration-specific macrophage signaling center for patterned bone formation and suggest that targeting the metabolic conditions that drive this population could improve the efficacy of regenerative therapies.

## INTRODUCTION

Bone is one of the few mammalian tissues capable of scarless repair, yet capacity is limited. Following fracture or amputation, bone regeneration requires precise spatial coordination where new bone must form in the right place, at the right time, and in the right amount. When this coordination fails, the consequences range from non-union and impaired regeneration to pathological mineralization of soft tissues^1–3^. Failed bone regeneration following large segmental defects, including total limb amputation, remains a significant clinical challenge, with current therapies relying on bone grafts, scaffolds, or exogenous growth factor delivery that promote bulk bone formation without the spatial precision needed for complete regeneration of function^4–7^. Together, these conditions highlight a gap in our understanding: we do not yet know how bone patterning is spatially controlled after injury. Understanding the cells and signals that instruct where bone is placed or not would allow us to target bone healing without side effects of mineralized soft tissues.

Macrophages are a critical but incompletely understood signaling center for osteogenesis during skeletal repair. After tibial fracture, macrophage depletion disrupts endochondral ossification by delaying callus formation and inhibiting mineralization^8–11^. After complete amputation, macrophage depletion stalls epidermal closure and prevents new bone and soft tissue growth^12,13^. In vitro, macrophages interact directly with osteoblasts and osteoprogenitor cells to promote their differentiation and mineralization^14–19^, while bone-associated macrophages, or osteomacs, have been shown to physically associate with bone-forming surfaces and support mineralization in vivo^9,16,20^. Critically, macrophages are also implicated in pathological bone formation. In heterotopic ossification, macrophage-derived signals in the early post-injury environment are shown to drive myoblast ossification of muscle tissue^21,22^, and macrophage depletion in some heterotopic ossification models reduces the amount of ectopic bone^23,24^. These observations raise the possibility that macrophages help specify where bone forms, acting as spatial regulators of mineralized tissue.

One mechanism by which macrophages could exert this spatial control is through the local secretion of factors including bone morphogenetic proteins (BMPs). BMPs are potent inducers of osteogenic differentiation and are among the most well-characterized regulators of bone patterning during development and repair^25,26^. Recombinant BMP2 and BMP7 have been in clinical use for over two decades to promote spinal fusion and long bone repair, yet outcomes remain variable, with complications including insufficient bone formation, ectopic ossification, and inflammation-driven resorption^26,27^. These clinical challenges reflect a broader biological reality: BMP signaling is highly context-dependent, shaped by the concentration, timing, and cellular environment in which BMPs are presented^28–31^. While osteoblasts, chondrocytes, and periosteal cells are established sources of BMP ligands during repair^32^, macrophages have also been shown to express and secrete BMP2 in vitro^14,19^. More recently, macrophage-derived BMPs have been linked to osteoinductive signaling following efferocytotic uptake of lipid-laden apoptotic cells, suggesting that macrophage metabolic state may regulate their osteogenic paracrine output^33^. Despite this, the identity of the specific macrophage population responsible for BMP secretion in vivo, its spatial relationship to sites of active bone formation, and the environmental signals that govern its activity during tissue repair remain poorly defined.

A potential explanation for this gap is that macrophages are not a uniform population within healing tissue. Single-cell transcriptomic studies have revealed extensive macrophage heterogeneity at sites of injury and repair, with distinct subpopulations defined by inflammatory state and tissue localization^34–36^ . We and others have shown that regenerative injuries are characterized by distinct macrophage transcriptional states compared to scar-forming injuries, and that these differences are driven at least in part by the intrinsic macrophage identity^37,38^. Within a given injury, macrophage subpopulations may occupy spatially distinct niches^39^ and engage in distinct paracrine interactions with local stromal cells^40^ potentially creating microenvironments that differentially support regeneration, repair, or scarring. If a specific macrophage subpopulation is responsible for BMP-mediated osteogenic patterning, it would represent a tractable cellular target for both promoting bone regeneration and suppressing pathological mineralization.

To test this idea, we make use of the mouse digit tip amputation model, which offers a unique opportunity to compare regenerative and scar-forming outcomes within the same animal under controlled conditions^41^. Amputation through the distal third phalangeal element (P3) results in complete regeneration of bone, dermis, epidermis, nerves, and vasculature through a blastema-mediated process^42–47^. In contrast, amputation just millimeters proximal through the second phalangeal element (P2) results in disorganized bone callus formation and dermal scarring, without restoration of patterned tissue^48,49^. Because both injuries occur in the same animal, differences in healing outcomes can be attributed to local rather than systemic factors, making this an ideal system for identifying the cellular and molecular determinants of regeneration versus scarring^50^.

Here, using integrated single-cell RNA sequencing, spatial transcriptomics, and in vitro metabolic analyses, we identify a spatially restricted inflammatory macrophage subpopulation that is unique to the regenerative injury environment, localizes to the growing bone front, and drives BMP-mediated osteogenic patterning through a fatty acid oxidation-dependent mechanism. This signaling center is unique to the regenerating bone and is absent in scar-forming tissue. This population and its unique metabolic and paracrine functions provide a key to understanding the divergence between regeneration and scar formation in bone injury.

## METHODS

### Animal care

All experiments were performed in accordance with the standard operating procedures approved by the Institutional Animal Care and Use Committee of LSU Health Sciences Center (Protocol #4754), Mayo Clinic (Protocol #A00007778-23) and University of Kentucky (Protocol #2024-4471).

### Amputations

For amputations, 8-week-old female CD1 (Charles River #022) and C57/Black6 (Charles River #027) mice were anesthetized with 3% isoflurane and maintained under anesthesia at 1% isoflurane for the duration of the procedure. Hind digits 2 and 4 were amputated at the P3 or P2 level as outlined in our previous publication^41^ and digit 3 was retained as an unamputated control. Digits were collected for analysis at various timepoints after amputation as indicated in the text.

### Immunofluorescence

Digits were collected at day 0, 13, and 17 and fixed in Neutral Buffered Formalin for 24 hours. Tissue was decalcified using Decal 1 (Leica 3800440) for 48 hours, washed with PBS and stored in 70% ethanol until paraffin processing and embedding. 4 µm sections were collected for staining. For staining, tissue was deparaffinized and incubated in heat retrieval solution (pH 6) overnight at 60° C. To block non-specific binding, slides were incubated in serum free blocking buffer (ScyTek AAA999). Staining was carried out at 4°C overnight using the following primary antibodies: pSMAD1/5 (Cell Signaling, Cat# 9516, Clone 41D10 0.1μg/mL), IBA1 (FujiFilm Cat# 011-27991; 1:1000), CD206 (R&D Systems Cat# AF2535; 1μg/mL), CD14 (Novus Biologicals Cat# NBP-51121; 5μg/mL). Slides were washed with PBST and incubated at room temperature for 45 minutes with secondary antibodies at 1:800 dilution in serum free block: donkey anti rabbit Alexa Fluor 568 (Invitrogen Cat# A10042), donkey anti goat Alexa Fluor 647 (Invitrogen Cat# A21447), and donkey anti rat AF488 (Invitrogen Cat#A21208).Nuclei were counterstained with DAPI (Thermo Fisher 3375) and slides coverslipped with ProLong Gold antifade reagent (Invitrogen Cat# P36930). Images were collected on an Olympus ix85 deconvolution microscope at 100x, 200x, or 600x from the center section of the digit. Representative images shown from an n=4 animals (n=8 digits).

### Bioinformatic analysis

Previously published scRNAseq data sets^51,52^ obtained from unamputated digits as well as day 7, 10 and 14 post amputation of P3 digits and day 10 and 14 post-amputation from P2 digits were downloaded from the Gene Expression Omnibus repository (GSE135985, GSE143888) and analyzed in *Seurat*. Cell clusters were identified using standard marker genes as we have previously published^53^. To calculate metabolic transcriptional profiles, the *AddModuleScore* function was used in combination with gene lists derived from metabolic pathways (Suppl Table 1). Variance calculations and statistical analyses were conducted using the ks.test run at default parameters from R package *stats* v4.4.0. Cell-cell communication between macrophages and fibroblasts was analyzed using NicheNet and CellChat^54,55^.

Previously published spatial transcriptomics data sets obtained from day 10 post amputation of P3 digits were downloaded from the Gene Expression Omnibus repository (GSE180682) and analyzed in *Seurat* ^53^. For mapping of macrophage phenotypes to spatial data, a gene signature was first generated for each single cell cluster by conducting differential gene expression relative to all other clusters. The top 50 DEGs were selected and combined with the *AddModuleScore* function to generate a predictive map of macrophage position. BMP signaling was assessed as described above for metabolic pathways using gene lists generated from KEGG pathways (Suppl Table 2).

### Metabolomics

To isolate macrophages, P2 and P3 amputated digits were collected at day 10 from 24 animals for a total of 48 digits (2 P3 amputations and 2 P2 amputations from each animal). The 48 digits were split into 3 groups for an n =3 technical replicates. To create a single cell suspension digits were treated both mechanically and enzymatically as previously described with modifications^56^. Briefly, tissue was digested with a 1:1 trypsin, dispase solution for 1 hr at 37°C allowing for subsequent mechanical separation of the epidermis. Separated epidermis and dermis plus bone were incubated with a solution of collagenase (1 mg/mL, VWR RLMB120-0100), hyaluronidase (0.5 mg/mL, VWR IC1512780), and elastase (0.015 U/μL, VWR IB1753-MC) in HBSS (VWR 45000–456) for 1 hr at 37°C. Following digestion, cell suspensions were washed with PBS and filtered through a 70 μm cell strainer. Single-cell suspensions were incubated with an FcγR block (CD16/32 block, 20 μg/mL, BD Pharmingen Cat# 553141) followed by incubation with directly conjugated primary antibodies at 4°C for 1 hr. Cells were collected if positive for CD53 and CD45 and negative for CD3. Antibodies included Alexa fluor 647 conjugated anti CD53 (BD Pharmingen Cat# 564931, PE-Cy7 conjugated anti CD45 (BD Pharmingen Cat# 561868), PE conjugated anti CD3 (BD Pharmingen 561799). Fluorescent-activated cell sorting (FACS) was carried out by trained experts in the Tulane University Flow Cytometry Core using the BD FACSAria Fusion sorter system (BD Biosciences, San Jose, CA). Laser calibration and compensation was performed for each experiment using unstained, single fluorescent, and fluorescent minus one (FMO) control samples. 3-5×104 isolated macrophages per group were snap frozen and shipped to Gigantest, Inc. (Baltimore, MD) for metabolic analysis.

Metabolites were extracted from the isolated macrophages using previously published protocols^57^. Metabolomics data was collected using a Thermo-Scientific IQ-X mass spectrometer with a Vanquish Horizon Binary ultra-performance liquid chromatography (UPLC) system. Data was analyzed using Thermo-Scientific XCalibur, Compound Discoverer, and TraceFinder software. Principal Components Analysis (PCA) and orthogonal partial least squares discriminant analysis (OPLS-DA) was used to characterize differential expression between groups^58–60^, identifying biomarkers with discriminatory power.

### BMDM isolation and culture

Macrophage progenitors were isolated from femur and tibia of C57BL/6NCrl (Charles River #027) or CD1 (Charles River #022) mice. After sacrifice, femur and tibia were surgically removed, and mechanically cleared of all skin, muscle and tendon. Marrow was aspirated from bones by flushing with 10 mL of DMEM (Corning 15-013-CV), 10% FBS through a 28-gauge syringe. Red blood cells were lysed with a hypotonic solution (Millipore Sigma Cat# 11814389001) and remaining cells were plated at a density of 8 × 10^6^ cells in T-75 culture flasks. For the first 7 days, bone marrow cells were grown in DMEM media (Corning 15-013-CV) supplemented with 20% L929 media containing M-CSF, 10% FBS, 1% PenStrep. After 7 days in culture, BMDM were split using cold PBS and a cell scraper, plated in 24-well plates at a density of 2 × 10^5^ cells/well and allowed to settle for 24 hr in DMEM media (Corning 15-013-CV) supplemented with 10% FBS and 1% PenStrep. For macrophage stimulation in high glucose (4.6g/L) conditions, cells were treated with increasing concentrations of palmitate-BSA (Agilent Cat#102720-100) or BSA alone in either high glucose culture media (DMEM Corning 15-013-CV, 10% FBS, 1% PenStrep). In low glucose conditions, macrophages were cultured in substrate limited media (DMEM Corning 17-207-CV, 0.5mM Glucose, 1.0mM glutaMAX, 0.5mM Carnitine, 1%FBS, 1% PenStrep) for 24 hours, then treated with various concentrations of palmitate-BSA or BSA in substrate enhanced media (DMEM Corning 17-207-CV, 5mM HEPES, 2.5mM glucose, 1.0mM glutaMAX, 0.5mM Carnitine, 1%FBS, 1% PenStrep) for 24 hours. Cells were then collected for qPCR or Seahorse analysis as below.

### RT-qPCR

4×10^5^ cells were collected for RNA isolation in 400uL of Trizol and RNA isolated via addition of BCP to separate phases and isopropanol to pellet the RNA according to manufacturer directions.

RNA pellets were washed in 70% EtOH twice and concentrations measured using a nanophotometer. 1ug of RNA was converted to cDNA using the high capacity cDNA kit (Applied Biosystems, Cat# 4368814). 4ng of cDNA were then anaylzed via quantitative PCR. qPCR using Powerup Sybr Green (Applied Biosystems Cat# A25741) was run on an Azure Cielo machine (Azure Biosytems, Dublin, CA) according to manufacturer directions with the following primers

Glut1 – CAGTTCGGCTATAACACTGGTG, GCCCCCGACAGAGAAGAT

Mdm2 – GGGAGTGATCTGAAGGATCC, CTCATCTGTGTTCTCTTCTGTC

Bmp2 - Mm.PT.58.10419414 (IDT premade)

Bmp4 - Mm.PT.58.28628134 (IDT premade)

Bmp5 - Mm.PT.58.10850304 (IDT premade)

Cpt1a – GCAAACTGGACCGAGAAGAG, CCTTGAAGAAGCGACCTTTG

### Seahorse Analysis

To create homogenate media, digits amputated at the P2 or P3 digits were isolated from the mouse 10 days after amputation. Digits were homogenized in DMEM (4 digits per mL of DMEM) using a handheld mechanical homogenizer. Samples were then filtered through a 70μm filter and stored at -80°C until use. Immediately prior to use homogenized samples were filtered through 0.22μm filters to sterilize for culture.

BMDM were harvested from tibia and femur and allowed to expand in a 100 mm plate before being reseeded at 80K cells/well in a 96-well Agilent Seahorse plate (Agilent, Santa Clara, CA). Cells were incubated with either P2 or P3 homogenate media for 24 hours and evaluated the following day using the Seahorse GlycoPER and Mito Stress Tests (Agilent, Santa Clara, CA) using the standard protocol and 0.75 μM FCCP concentration. (*N* = 7-10 wells per group) and as previously described^53^. To test FAO, macrophages were prepared as and seeded as described above. Cells were changed to P2 or P3 media for 24 hours and evaluated the following day using the Seahorse XF Palmitate Oxidation Stress Test using the standard protocol with an acute etomoxir injection. For palmitate supplementation, BMDM were plated in seahorse plates with either high glucose culture media (DMEM Corning 15-013-CV, 10% FBS, 1% PenStrep) or low glucose culture medium (DMEM Corning #17-207-CV, 0.5mM Glucose, 1.0mM glutaMAX, 0.5mM Carnitine, 1%FBS, 1% PenStrep) for 24 hours. In high glucose conditions, palmitate-BSA or BSA alone (200nM) was added to fresh high glucose culture media and cells incubated for 1 hour prior to Seahorse analysis. For low glucose conditions, palmitate-BSA or BSA alone (200nM) was added to substrate enhanced media (DMEM Corning #17-207-CV, 5mM HEPES, 2.5mM glucose, 1.0mM glutaMAX, 0.5mM Carnitine, 1%FBS, 1% PenStrep) for 1 hour prior to Seahorse analysis.

## RESULTS

### Unique transcriptional profiles of inflammatory and resolution macrophage subtypes between P2 and P3 injury

We first re-analyzed single-cell RNA sequencing (scRNAseq) data sets from the regenerating (P3) and fibrotic (P2) digits^51,52^ to identify cellular subtypes in both injuries, as well as in uninjured conditions (Fig. 1A, Suppl Fig. 1A). Analyses identified five different clusters of myeloid lineage cells defined by expression of *Cd53*, *Fcer1g*^61^ and lack of expression of T-cell marker *Cd3*(clusters 8, 9, 10, 11, and 12 - Fig. 1B, Suppl Fig. 1B). Monocytes (cluster 8) showed enriched expression in *Plac8* and *Ccr2* (Fig. 1C). Inflammatory macrophages (cluster 9) expressed elevated levels of chemokine and chemokine receptor genes (*Ccl3, Ccl4, Cxcl2, Cxcl3*), calcium-binding proteins (*S100a8, S100a9*), enzymes and metabolic regulators (*Irg1, Hdc, Upp1, Ptgs2*) and proinflammatory genes (*Il1a*, *Il1b*, *Il1rn*, *Slpi*, *Marcksl1*, *Isg15*) (Fig. 1C). Resolution macrophages (cluster 10) were identified via expression of specific surface receptors known to trigger anti-inflammatory responses (*Mrc1, Trem2, Csf1r, Pf4*), genes involved in antimicrobial defense (*Lyz2, Cd36, Npl, Lgmn, Ctsb, Syngr1*), high expression of complement proteins (*C1qc, C1qa, C1qb*) and the selenoprotein, *Selenop*. Dendritic cells (cluster 11) showed high expression of genes involved in antigen presentation and processing (H*2-Ab1, H2-Aa, H2-Eb1, H2-DMa, H2-DMb1, Cd74, Batf3*) as well as the dendritic cell specific antigen and chemokines receptors *Cd209a* and *Ccr7*. Finally, osteoclasts (cluster 12) were identified based on expression of genes related to bone resorption, including proteolytic degradation of bone matrix (*Ctsk, Mmp9*), acidification of the resorption lacunae (*Atp6v0d2, Atp6v1b2*), regulation of osteoclast differentiation (*Nfatc1*), and mitochondrial metabolism (*Uqcr10, Uqcr11*) (Fig. 1C). Each cell cluster was present at each time point (Fig. 1D) and in both injury types (Fig. 1E), however the ratios of each cell cluster differed between injuries (Fig. 1F).

**Figure 1.**
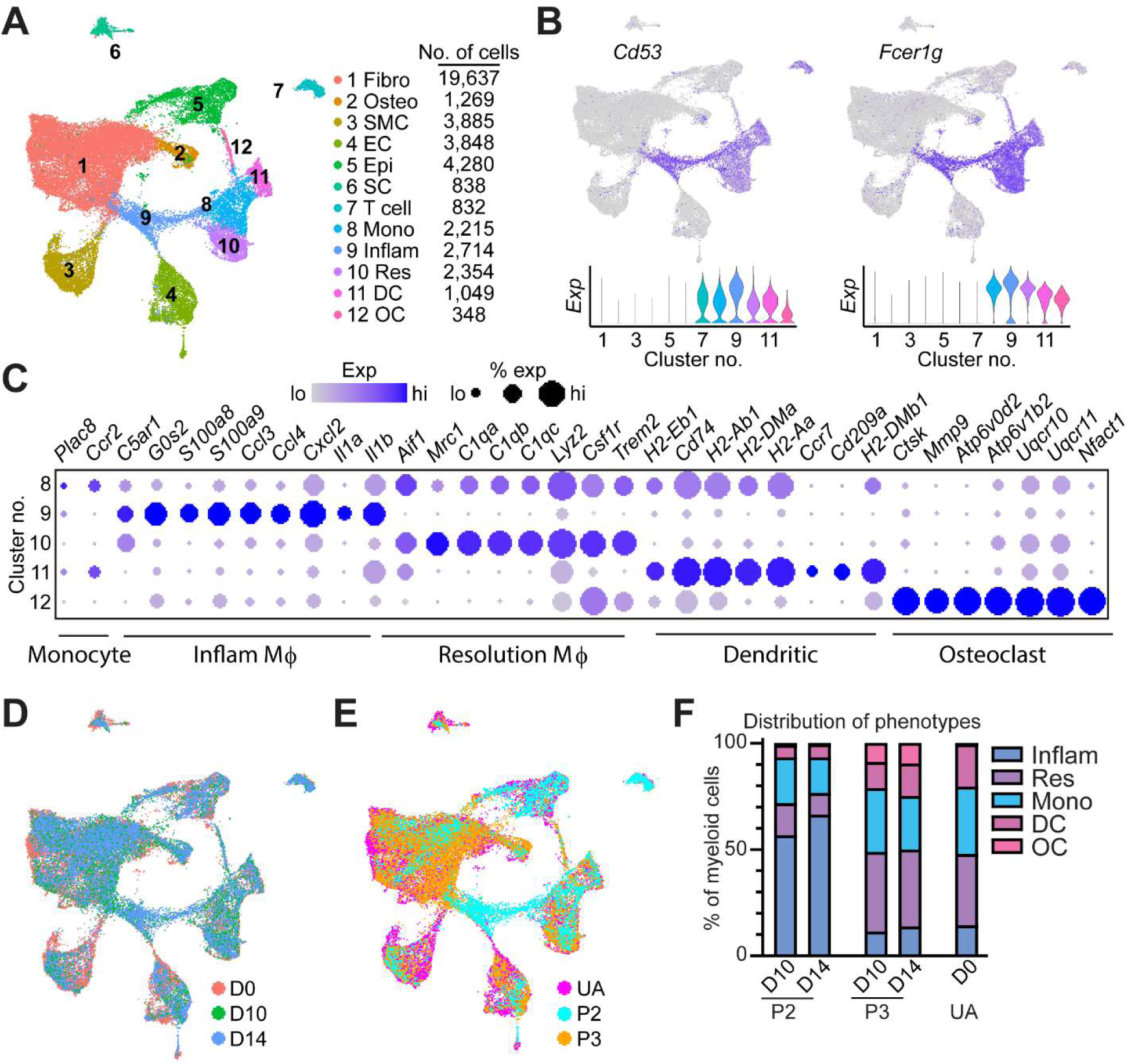
ScRNAseq reveals the prolonged presence of inflammatory macrophages following fibrotic injuries. A) UMAP of all cells from P2 (non-regenerative) and P3 (regenerative) amputations at three timepoints: Day 0, D10 and D14. B) Feature (top) and violin plots (bottom) showing expression of the hematopoietic marker *Cd53*, and the myeloid cell marker *Fcer1g* used to identify myeloid populations (clusters 8-12). C) Dot plot showing marker gene expression used to distinguish myeloid lineage cell populations as monocytes (cluster 8), inflammatory macrophages (cluster 9), resolution macrophages (cluster 10), dendritic cells (cluster 11) and osteoclasts (cluster 12). D) UMAP colored by time point to distinguish contribution to each cluster based on time. E) UMAP colored by injury type to distinguish contribution to each cluster based on injury type. UA, unamputated. F) Quantification showing relative contribution of each myeloid lineage cell type across each injury type and time point.

An analysis of the inflammatory macrophage cluster (cluster 9) over time shows that at day 7 post amputation (D7) inflammatory macrophages spike in P3, rising to constitute 43% of the population (Suppl Fig. 1C) before decreasing at D10. While the overall abundance was higher, the relative ratios of cells within the myeloid lineage at D10 and D14 following P3 injury was similar to that of uninjured digits, except for an expanded osteoclast group (Fig. 1F). In contrast, an analysis of the same cluster in P2 shows a notable expansion of inflammatory macrophages at D10 that was sustained through D14. This increase in P2 inflammatory macrophages was accompanied by a relative decrease in monocyte and resolution macrophage populations (Fig. 1F). These data suggest that while both P2 and P3 injuries induce inflammatory macrophage expansion, this induction persists during P2 fibrotic scarring but not during P3 regeneration. Further, these data support a P3 myeloid cell abundance profile that is closer to that of the uninjured digit during blastema formation and regrowth phases (Day 10 and Day 14).

Recent cross-species comparison studies suggest that inflammatory macrophages, defined by expression of pro-inflammatory cytokines such as *Tnf*, *Il6*, and *Il1b* as well as surface markers such as *S100a8/9* and *Cd14*, are transcriptionally distinct when derived from regenerative vs scar forming injuries.^37^ However, these findings do not take into account species-specific immune system differences. In light of this, we next compared inflammatory and resolution macrophage transcriptomes from fibrotic P2 and regenerative P3 injuries of the mouse digit. We isolated and re-analyzed monocyte, inflammatory and resolution macrophage clusters (clusters, 8, 9, and 10) from fibrotic and regenerative injuries (Fig. 2A, B). Surprisingly, we found that the variance in gene expression across myeloid cells was accounted for more by injury type, rather than cluster (Fig. 2C), indicating that transcriptional changes are driven predominantly by injury type rather than monocyte/macrophage subtype. Additionally, inflammatory macrophages showed the highest transcriptional variability between injury types, followed by resolution macrophages and then monocytes (Fig 2D) which indicates changes in inflammatory macrophages may have a greater contribution to changes in healing outcomes.

**Figure 2.**
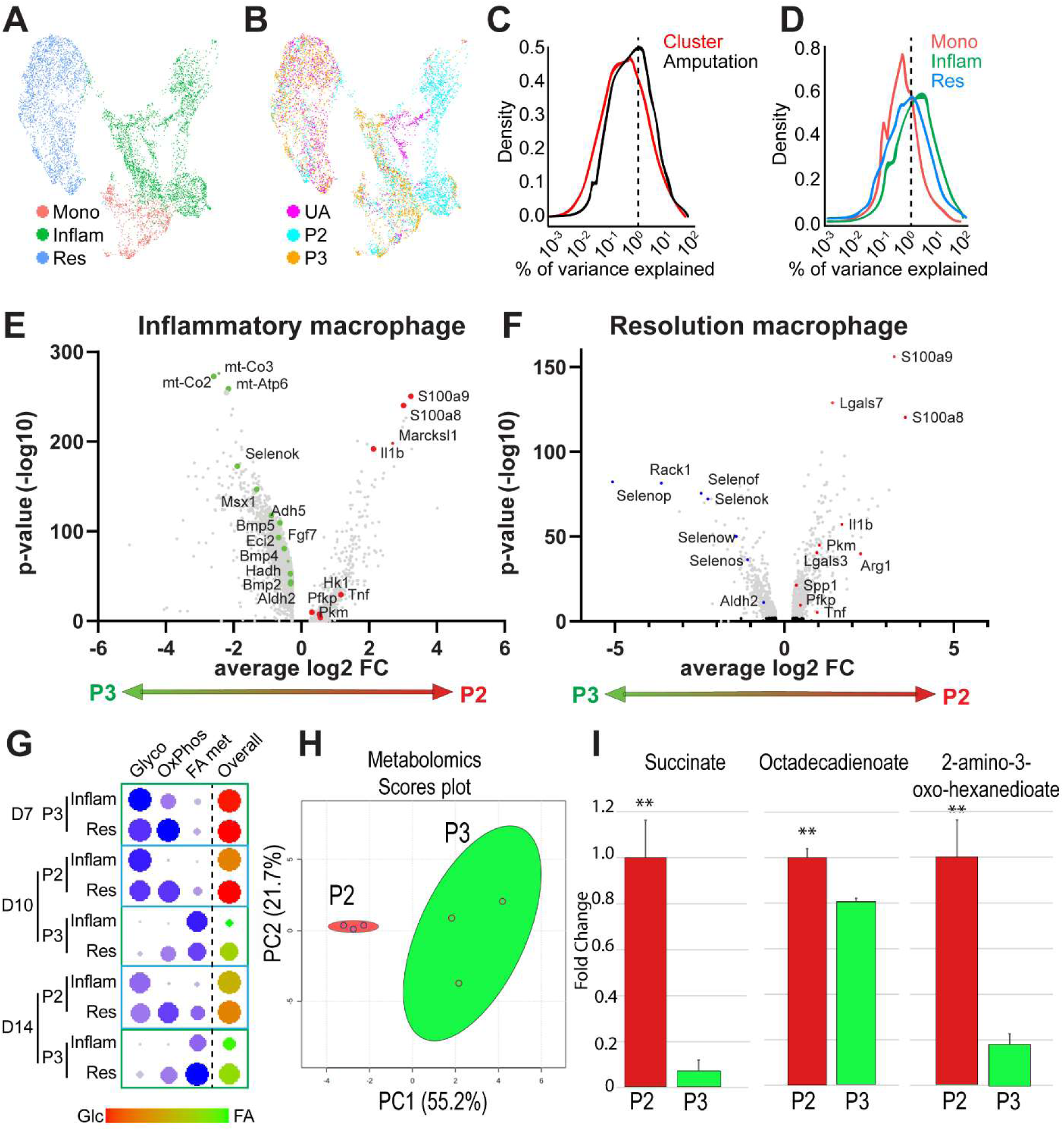
Macrophages show injury type-dependent metabolic shifts in glucose and fatty acid metabolism. A) UMAP of monocyte and macrophage clusters (clusters 8-10 from Fig 1). B) UMAP colored by injury type. UA = unamputated, P2 = non regenerative, P3 = regenerative. C) Analysis comparing variance in gene expression accounted for by injury type or clusters. D) Analysis comparing variance in gene expression accounted for by individual cell type. E) Volcano plot of differentially expressed genes in the inflammatory macrophage cluster comparing P3 (green) versus P2 (red) injuries (all time points). F) Volcano plot of differentially expressed genes in the resolution macrophage cluster comparing P3 and P2 injuries (all time points). G) Analysis of genes involved in the glycolytic (glyco), oxidative phosphorylation (oxphos), and fatty acid metabolism (FA met) metabolic pathways. Overall score indicates the ratio between glycolysis (glc, red) and fatty acid metabolism (green) in each cluster. Inflam (inflammatory macrophage), Res (resolution macrophage)D = day post amputation. H) PCA of metabolomic data obtained from D10 macrophages following P2 or P3 amputation. I) Quantification of metabolite differences between P2 and P3 macrophages. Succinate and fatty acid-related metabolites, Octadecadienoate and 2-amino-3-oxo-hexanedioate. n=3, ** p< 0.01

We next investigated differentially expressed genes (DEGs) within inflammatory and resolution macrophage subtypes in P2 and P3 injury. Volcano plot analysis of inflammatory macrophages at D10 and D14 showed upregulation of enzymes involved in glycolysis (*Hk1*, *Pfkp*, *Pkm*), and inflammation pathways: NFκB signaling (e.g. *Cxcl3*, *Cxcl2*, *Icam1*, *Ccl4*, *Cd14, S100a8, S100a9*), TNF signaling (e.g. *Tnf*, *Map2k3*, *Cebpb*, *Il15*, *Traf1*, *Ptgs2*) and chemokine signaling (e.g. *Il1b*, *Stat1*, *Ccr1*, *Cxcr4*, *Ccl6*) in P2 inflammatory macrophages. In contrast, P3 inflammatory macrophages showed a significant upregulation of genes involved in fatty acid metabolism (e.g. *Acaa2*, *Acat1*, *Adh5*, *Aldh2*, *Echs1*, *Eci2*, *Hadh*, *Hadha*), oxidative phosphorylation (e.g. *mt-Co2*, *mt-Co3*, *mt-Atp6*), selenoproteins (e.g. *Selenow*, *Selenos*, *Selenok*, *Selenof*, *Selenop*) and growth factors (e.g. *Bmp2*, *Bmp4, Bmp5*, *Fgf7*) (Fig. 2E, Suppl Table 3). A parallel volcano plot analysis of P2 and P3 resolution macrophages (Fig. 2F) showed upregulation of genes involved in fibrosis and repair (*Arg1*, *Lgals3*, *Spp1*) in P2 versus P3. In contrast, P3 resolution macrophages showed upregulation of genes involved in fatty acid oxidation (e.g. *Aldh2*) and selenoproteins that regulate oxidative stress (*SelenoP*, *SelenoS*, *SelenoK, SelenoW*)^62–65^, a profile similar to P3 inflammatory macrophages (Fig.2F, Suppl Table 4).

To further investigate these deviations in metabolic profiles occurring within P2 and P3 macrophages, we next ran module scores on gene lists to get a more global view of the metabolic profile of these cells. This function calculates the average expression of a list of 20 -100 genes from established KEGG pathways and published sources related to each metabolic process and creates a single output, normalized to background, to quantify the combined transcriptional regulation of each metabolic profile (Suppl Table 1). Using this analysis, we found that both inflammatory and resolution P3 macrophages prefer glucose-dependent metabolic pathways (glycolysis and oxidative phosphorylation) during the osteoclast-driven histolytic phase (D7) (Fig 2G). This glycolytic preference shifts at D10 in P3 macrophages to FAO-associated metabolic genes, while P2 macrophages of both subtypes continue to utilize glycolysis and oxidative phosphorylation (Fig. 2G), highlighting a clear metabolic difference at a critical regenerative time point that persists through D14.

To confirm these findings, we conducted metabolomics on FACS-isolated (CD45^+^/CD53^+^/CD3^-^) macrophages from both injury types at D10. PCA analysis confirmed a unique metabolic signature between P2 and P3 macrophages (Fig. 2H). Supporting our findings that P3 inflammatory macrophages preferentially utilize FAO, P3 inflammatory macrophages showed decreased levels of fatty acid-related metabolites, Octadecadienoate and 2-amino-3-oxo-hexanedioate, whereas P2 inflammatory macrophages showed increased levels of the FAO inhibitor and glycolysis by-product, succinate (Fig. 2I). Together these data suggest both inflammatory and resolution macrophage subtypes are transcriptionally and metabolically distinct between regenerating and scar forming injuries, where the regenerative metabolic profile is defined by a shift from glucose metabolism (D7) to FAO (D10-14) with increased utilization of long-chain fatty acids.

### Environmental factors in the injury environment drive changes in macrophage cell metabolism

To test whether signals from the injury environment dictate metabolic differences seen in P2 and P3 macrophages in vivo, we cultured bone marrow derived macrophages (BMDM) in media containing homogenized injured tissue collected at D10 from P2 and P3 digit amputations (Fig. 3A). We then used Seahorse analysis to measure changes in glycolytic rate (GlycoPER, Glycolytic Rate Assay) and oxygen consumption rate (OCR, Mito Stress Test) in tissue homogenate treated BMDMs (Fig. 3, Suppl Fig. 2A-C). Glycolytic rate analysis showed increased GlycoPER in cells exposed to both P2 and P3 homogenate, with significantly higher rates observed in P2 homogenate-exposed cells (Fig. 3B). In parallel, basal OCR was decreased in BMDM pre-treated with either P2 or P3 homogenate compared to control BMDM with no significant difference in maximal respiration rates between groups (Fig. 3C) suggesting a reliance on non-mitochondrial sources of energy when treated with P3 or P2 homogenate. Treatment with the FAO inhibitor etomoxir (Eto) to prevent the use of lipids as a substrate showed a decrease in OCR in control and P3-treated groups but not in P2-treated groups, suggesting cells exposed to P2 homogenate media are less reliant on FAO than cells exposed to control or P3 homogenate (Fig. 3D). Together these data indicate that macrophage metabolism can be directed by the local injury environment, with P3 promoting reliance on FAO and glycolysis and P2 promoting a reliance primarily on glycolysis.

**Figure 3.**
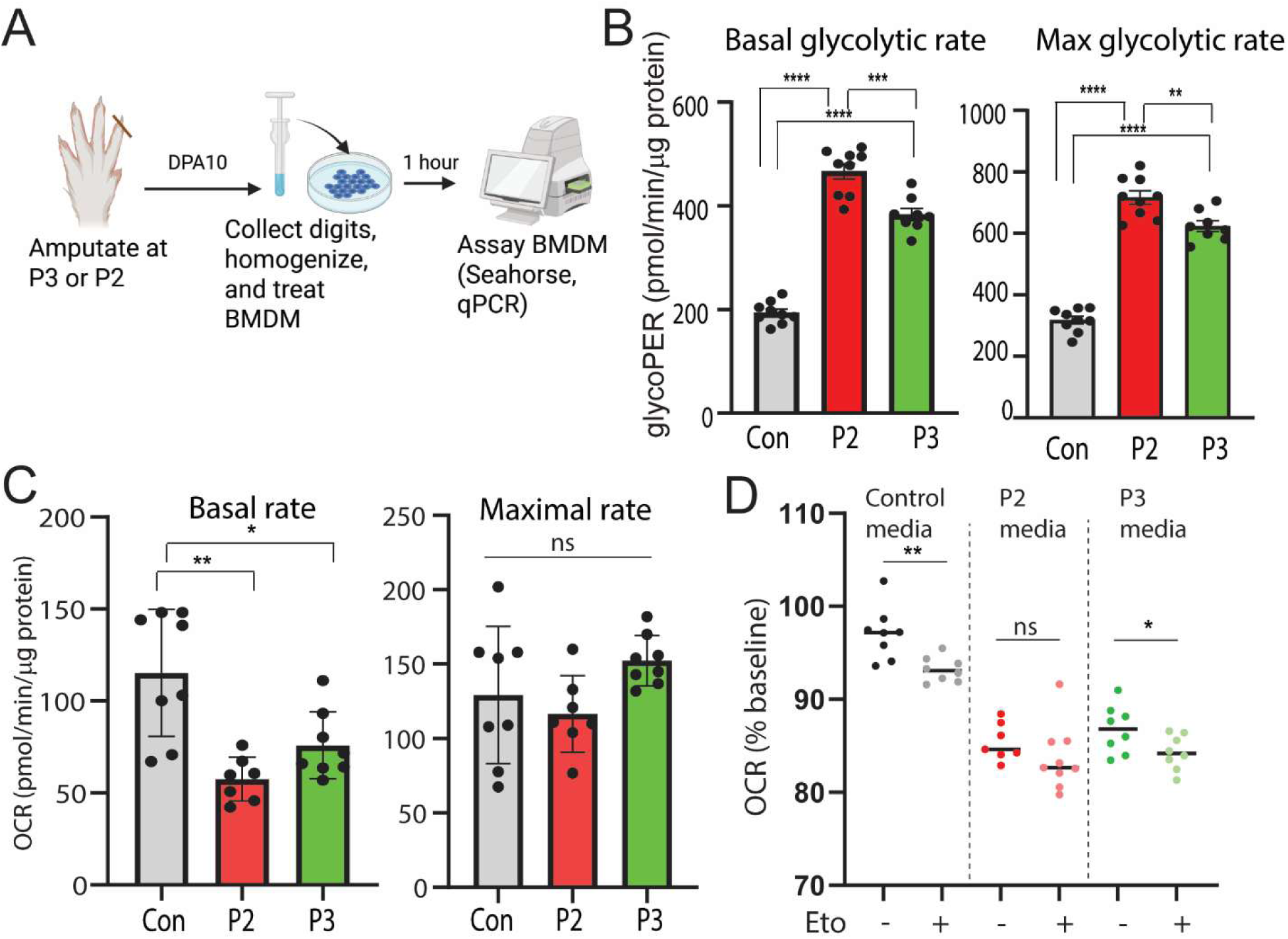
Differential metabolic reprogramming of bone marrow derived macrophages by P3 or P2 injury environments. A) Schematic of the experimental design. Digits are amputated at the P2 or P3 level and collected 10 days post amputation (DPA). Tissue from the amputation is homogenized and BMDM is exposed to the homogenized tissue for 1 hour. BMDM metabolic rates are then analyzed via Seahorse analysis. B) Seahorse Glycolytic Rate analysis in BMDM exposed to control media, P2 homogenate or P3 homogenate. Comparison of basal and maximal glycolytic proton efflux rate (GlycoPER). Welch’s T-test *p<0.05, **p<0.01, ***p<0.001, ****p<0.0001, n=8-10. C) Seahorse Mito Stress Test in BMDM exposed to control media, P2 homogenate or P3 homogenate. Comparison of basal and maximal oxygen consumption rates (OCR) Welch’s T-test, *p<0.05, **p<0.01 n = 8-10. D) XF Palmitate Oxidation Stress Test analysis of FAO. BMDM are exposed to control media, P2 homogenate or P3 homogenate. An acute injection of etomoxir (Eto +) or saline (Eto -) is injected after 14 minutes of analysis. Data is graphed as percent change in basal OCR rate after Eto or saline injections. Welch’s T-test, *p<0.05, **p<0.01 n = 8-10.

### Inflammatory macrophages are associated with increased BMP signaling within the proximal blastema of regenerating P3

Our initial scRNAseq analysis identified several differences in expression of genes linked to metabolic, growth factor and cytokine signaling between P2 and P3 macrophages. To further understand the potential impact of these altered gene expression profiles within macrophages on the healing outcomes driven by the regenerative blastema fibroblasts, we conducted cell-cell communication analyses between macrophages and fibroblast cell types. Top predicted ligands derived from pooled (P2 and P3) monocyte/macrophage populations were consistent with our differential gene expression analysis (Fig. 2E, Fig. 1E,F) and included growth factors (*Fgf2*, *Bmp4*, *Tgfb1*) and the inflammatory cytokine *Tnf* (Fig. 4A). Additional analyses identified the cognate fibroblast receptors, as well as potential downstream targets regulated by these macrophage-derived ligands (Fig. 4B, C). Differential expression analyses for P2 and P3 inflammatory and resolution macrophages identified *Tnf* and *Bmp4* as top regulatory ligands enriched within P2 and P3 injuries, respectively, and derived primarily from inflammatory macrophages (Fig. 4D, E). To better understand how cells might be communicating with each other, we used cell chat analysis with a focus on BMP signaling. BMP ligands were most highly expressed in P3 inflammatory macrophages (cluster 9) and fibroblasts (cluster 1) with receptors present on fibroblasts, osteoblasts, smooth muscle cells, epithelial cells and schwann cells (clusters 1, 2, 3, 5, and 6 respectively) (Fig. 4F). In contrast, no BMP ligand-receptor signaling was predicted in P2.

**Figure 4.**
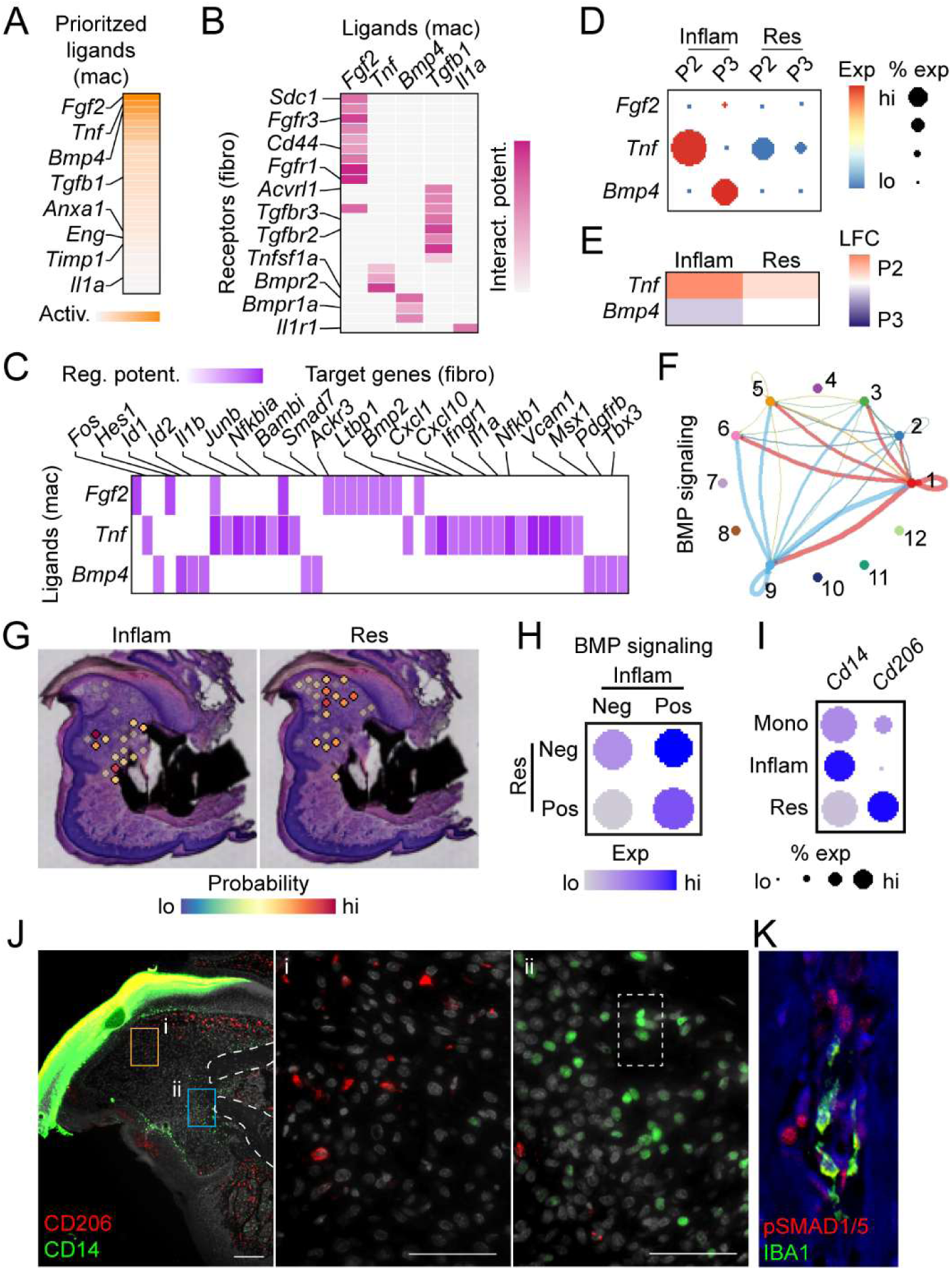
P3 inflammatory macrophages are spatially restricted and interact with fibroblasts through BMP signaling. A) Highest predicted ligand expression in P2 and P3 macrophage clusters (inflammatory and resolution subtypes combined). B) P2 and P3 ligand/receptor (macrophage/fibroblast) expression likelihood. C) Downstream targets of ligands identified in 3A and B in fibroblasts. D) P2 and P3 macrophage expression of top ligands by injury type. E) Difference in gene expression between P2 and P3 (red P2>P3, blue P2<P3). F) Circus plot confirming strong BMP signaling between inflammatory P3 macrophages (cluster 9) and fibroblasts (cluster 1). No signaling is predicted in P2. G) Localization of macrophage types in vivo using combined analysis of spatial transcriptomics and single cell RNAseq. H) Predicted BMP signaling as a function of inflammatory or resolution macrophage presence. I) Confirmation of macrophage subtype marker transcript expression of *Cd14* and *Cd206* at D10 in P3. J) Immunofluorescent staining of CD206 (resolution, red, inset (i)) and CD14 (inflammatory, green, inset (ii)) at D10 in P3. DAPI (grey) Scale bar = 100μm, n=4 digits. K) Immunofluorescent staining of pSMAD1/5 (red) and IBA1 (green) at D13. DAPI (blue) Scale bar = 100μm, n=4 digits

To localize inflammatory and resolution P3 macrophages and to determine the impact of the predicted BMP signaling on neighboring cells in vivo, we shifted from analysis of dissociated macrophages analyzed in scRNAseq to the use of spatial transcriptomic data from a P3 regenerating digit at D10 post-amputation. We used spatial deconvolution to determine the probability that an inflammatory or resolution macrophage single cell cluster was present within each spatial spot (Fig. 4G). These data show P3 inflammatory macrophages are restricted to near the bone stump, while the P3 resolution macrophages were found primarily in the more dorsal and distal area of the blastema. Next, spots were characterized as being either positive or negative for each of the macrophage populations based on predictive modeling values and analyzed for the presence of BMP activating transcripts (Fig. 4H). These results show that spatial spots predicted to contain inflammatory macrophages, but not resolution macrophages, showed the highest levels of BMP signaling transcripts. In contrast, spots devoid of inflammatory macrophages correlated with the lowest levels of BMP transcripts. To confirm these predictive transcriptional results, we next carried out immunofluorescent staining. Within our scRNAseq data, we identified *Cd14* and *Cd206* as markers for inflammatory and resolution macrophages, respectively (Fig. 4I). Immunostaining for these markers corroborated our spatial predictive modeling. CD14^+^ inflammatory macrophages localize near the bone stump at D10 while CD206^+^ resolution macrophages localize to the distal and ventral areas of the blastema (Fig. 4J).

Additionally, co-staining for phosphoSMAD1/5 (the downstream BMP effector proteins) and the pan macrophage marker IBA1 shows IBA1^+^ cells within close proximity to pSMAD1/5^+^ cells near growing bone in P3 amputations at D10, D14 and D17 after amputation (Fig. 4K, Suppl Fig. 3). Taken together these data support a specific role for P3 inflammatory macrophages in driving BMP-mediated tissue patterning and differentiation during regeneration.

### FAO, without an increase in glycolysis, drives BMP ligand expression in macrophages

In vivo, scRNAseq data showed an increase in FAO and *Bmp* ligand expression concurrent with a decrease in glycolysis in P3 macrophages (Fig. 2). Contrary to our in vivo data, BMDM exposed to homogenized tissue from either injury site did not demonstrate an increase in *Bmp2, 4, or 5* expression (Suppl Fig. 2D). However, observation of decreased fatty acid-related metabolites in vivo (Fig. 2I) suggest P3 cells may be responding to environmental fatty acids. To further explore the relationship between *Bmp* expression and metabolism, we exposed BMDM to the long chain fatty acid palmitate, with or without glucose supplementation, to create a dependency on FAO (Suppl Fig. 4A). When BMDM are exposed to palmitate in high glucose media, basal and maximal respiratory rates increase (Suppl Fig. 4B). Inhibition of FAO using etomoxir (Eto) to prevent the use of palmitate as a substrate showed a decrease in OCR, suggesting part of the palmitate-dependent increase in OCR is a reliance on beta oxidation (Suppl Fig. 4B). BMDM in these palmitate and glucose-supplemented conditions also display an increase in compensatory glycolysis with no change in *Bmp* expression (Suppl Fig. 4C,D), similar to P3 homogenate-exposed BMDMs (Suppl. Fig. 2). However, when BMDM are exposed to palmitate without glucose supplementation, there is no significant change in the glycolytic rate (Fig. 5A) while the maximal OCR rate is reduced after exposure to Eto (Fig. 5B), suggesting a reliance on FAO and not glycolysis. These metabolic shifts correspond to an increase in both *Bmp2* and *5* expression in cells exposed to 20μM of palmitate with restricted glucose availability (Fig. 5C) and a simultaneous decrease in the glycolytic enzyme *Glut1* (Fig. 5C). Increasing concentrations of palmitate increased expression of *Glut1* while simultaneously decreasing expression of the *Bmps* (Fig. 5C). These data parallel our in vivo data showing that a palmitate-dependent increase in OCR, with no concurrent increase in glycolysis, promotes BMP expression in macrophages.

**Figure 5.**
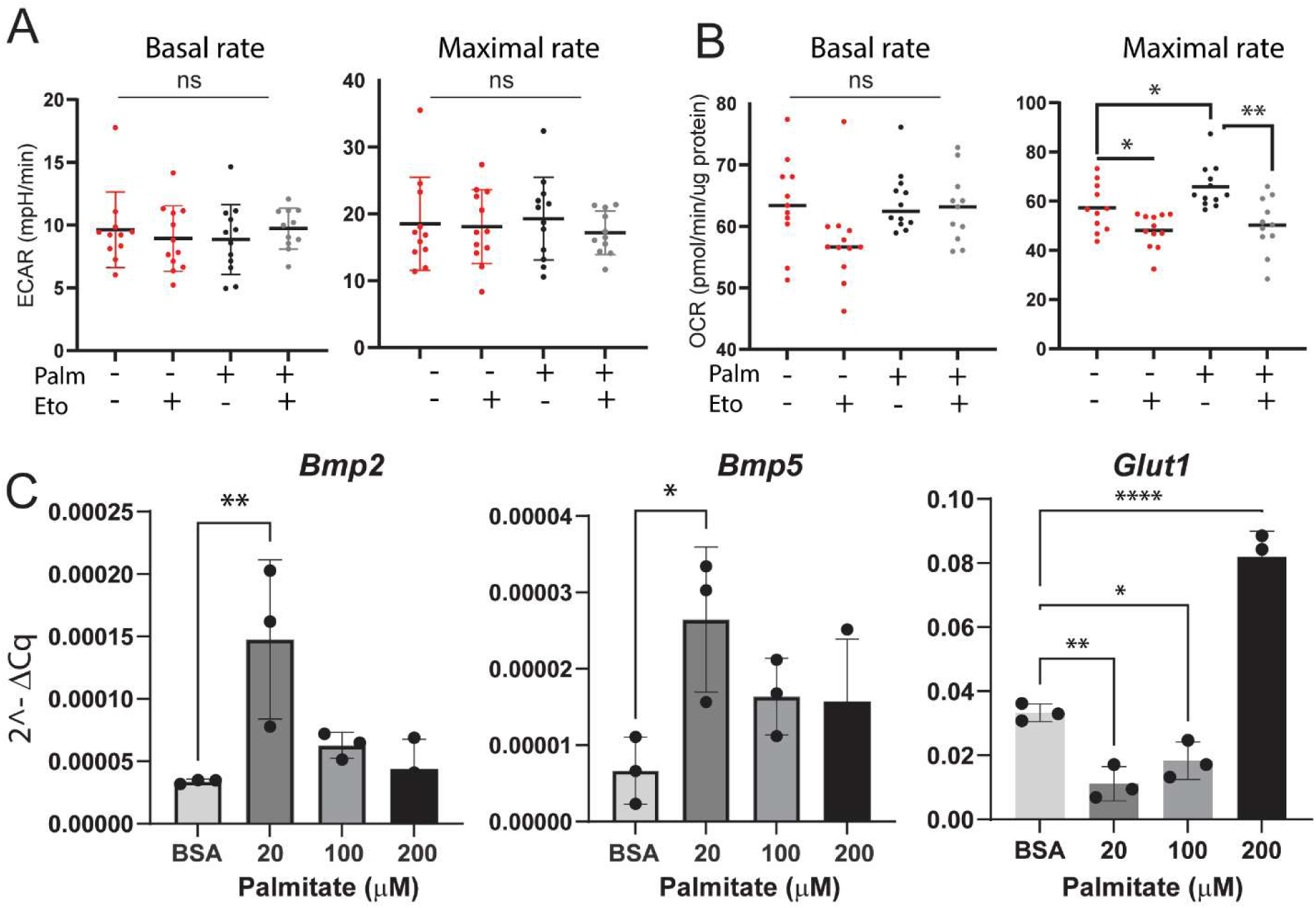
Increase in FAO without an increase in glycolysis drives BMP ligand expression in macrophages. Seahorse Palmitate Oxidation Stress Test analysis of FAO. BMDM were exposed to palmitate or BSA control in a low glucose environment. Seahorse analysis for ECAR (A) and OCR (B) basal and maximal respiratory rates. Welch’s T-test *p<0.05, **p<0.01, ***p<0.001, ****p<0.0001 C) Expression of *Bmp2, 4 and 5, and Glut1* in BMDM exposed to exposed to palmitate in low glucose media graphed as 2^ΔCq normalized to the housekeeping gene B2M. n=3 Dunnet’s multiple comparison test *p<0.05, **p<0.01, ***p<0.001, ****p<0.0001

## DISCUSSION

In this study, we identify a spatially restricted inflammatory macrophage subpopulation that functions as a signaling center for patterned bone regeneration. The population is present at the growing bone front in the regenerative injury and absent in scar-forming tissue. Using integrated single-cell RNA sequencing, spatial transcriptomics, and in vitro metabolic analyses in the mouse digit tip amputation model, we show that this population is defined by a specific transcriptional and metabolic identity, expresses BMP ligands that activate neighboring osteoblasts, and is governed by a two-part environmental metabolic switch involving increased fatty acid oxidation coupled with reduced glycolytic activity. These findings reframe our understanding of macrophage function in bone repair: rather than acting as broadly permissive or suppressive regulators of healing, a specific inflammatory macrophage subtype occupies a defined spatial niche and delivers precise osteogenic patterning cues at a critical window of regeneration. Critically, despite the presence of inflammatory macrophages in both injury types, only the P3 regenerative environment produces the conditions necessary for the emergence of this macrophage state.

Understanding how and to what degree cells change their wound healing microenvironment, or alternatively how the wound healing microenvironment changes cells, is paramount to being able to promote regeneration. Exposure to two different injury sites in close proximity leads to different metabolic reprogramming in macrophages. Our single-cell RNA-seq and spatial transcriptomic analyses identify two transcriptionally distinct clusters of activated macrophages within the digit injury site. Both clusters are present across all time points, but their proportions and metabolic signatures diverge between regenerative and non-regenerative injuries. In the fibrotic P2 injury, we observe a prolonged dominance of macrophages with high glycolytic gene expression and elevated pro-inflammatory mediators such as *Tnfα*, consistent with a chronic inflammatory environment ^66–75^ that resolves via scar formation. By contrast, in the regenerative P3 injury, both macrophage clusters shift away from glycolysis toward an FAO-enriched state consistent with anti-inflammatory macrophage identity ^76–78^ and suggesting that cues in the regenerative environment actively reprogram macrophage metabolism, independent of baseline cluster identity. Similarly, our ex vivo stimulation experiments support the idea that the metabolic differences seen in P2 and P3 inflammatory macrophages are driven by inherent differences in the injury microenvironment rather than intrinsic differences in each macrophage subtype. When bone marrow derived macrophages are exposed to homogenized P3 tissue, they upregulate FAO-associated pathways, whereas P2 homogenates preferentially induce a glycolytic program. These data indicate that regeneration-competent tissue contains factors that promote a FAO-biased metabolic state in macrophages, while scar-forming tissue reinforces glycolytic, pro-inflammatory metabolism^66,67,74^. This environmentally driven metabolic divergence raises a direct question: what is it about the FAO-enriched state that supports regeneration, and specifically, does it alter what macrophages communicate to surrounding cells?

Recent work has started to fill this gap by showing that the efferocytotic uptake of lipids from apoptotic cells can drive FAO in macrophages and, in turn, enhance their osteoinductive capacity through BMP production ^33^. Similarly, efferocytosis-driven FAO has been shown to couple mitochondrial metabolism to anti-inflammatory and tissue-repair gene expression programs^79^. Our data extend these findings in two important ways. First, we identify a BMP-expressing macrophage population in vivo within a regenerative bone injury, directly linking high FAO levels in macrophages to BMP ligand production in a model of epimorphic regeneration. Second, by comparing regenerative and non-regenerative contexts, we show that an increase in FAO is not sufficient on its own. Robust BMP expression is associated with a metabolic state characterized by elevated FAO accompanied by a relative reduction (or at least no further increase) in glycolytic activity. In contrast, when glycolysis remains high, macrophages maintain a pro-inflammatory profile and fail to initiate the BMP program even in the presence of exogenous fatty acids. The idea that macrophage-dependent regeneration is not just an either/or phenomenon, but a two-part change involving both a decrease in glycolysis *and* an increase in FAO paves the way to better understanding how these two metabolic processes in macrophages facilitate macrophage-osteoblast crosstalk through BMP signaling.

BMP is a potent regulator of bone formation, and recombinant human BMPs have been approved as clinical therapies for over 20 years^26^. Exogenous BMP2 can drive robust bone growth for spinal fusion, rescue non-unions, and induce regeneration in otherwise non-regenerative contexts, including digit and limb segments, in both neonatal and adult mouse models^25,28,80^. BMP expression defines the blastema, and periosteal responsiveness to BMP2 is a key determinant of whether a given amputation level can regenerate^28,53,81,82^. Despite this, clinical BMP therapies show variable success, with complications ranging from insufficient bone formation to excessive or ectopic bone^27^. Research supports that BMP outcomes are highly context dependent, shaped by surrounding factors in the tissue environment and the timing, concentration and location of BMP exposure relative to osteoprogenitors^27–31,83^. Our findings add to this framework by identifying an inflammatory macrophage population as an endogenous source of BMP ligands during the critical window of bone rebuilding (days 10–14 post-injury) and identifying environmental factors that modulate this population. Our spatial transcriptomic and immunofluorescence data reveal that these BMP-expressing macrophages are not broadly distributed across the injury site but are instead physically confined to the region immediately adjacent to the bone stump where new bone patterning is initiated. This spatial restriction highlights the idea that endogenous BMP delivery during regeneration is not simply a matter of the right signal at the right time, but also from cells positioned at the right place. In contrast, in the scar-forming P2 injury, macrophages fail to mount an equivalent BMP program and do not occupy osteogenic niches at the amputation plane. That this spatially restricted BMP-expressing population is entirely absent in the non-regenerative P2 injury suggests that the failure of patterned bone regeneration in scarring contexts may reflect not just a deficit in BMP availability, but a failure to establish the localized cellular source that delivers it precisely where it is needed.

Together, these findings support a model in which cues in the injury environment shift macrophage metabolic programming in spatially restricted areas to promote local cell-cell communication for patterned bone formation. In the regenerative digit tip, environmental cues (including fatty acids) bias macrophages toward a FAO-dominant state that enables BMP ligand production supporting organized bone regeneration. In the non-regenerative injury, persistent glycolytic metabolism reinforces inflammatory cytokine production, limits the emergence of the BMP-expressing macrophage state, and culminates in scar formation. These insights suggest that strategies capable of expanding or enhancing this population in a spatially and temporally controlled manner could improve the precision of bone growth after injury.

## Acknowledgements

This publication was supported by the NIH National Institute of Child Health and Human Development through grant number R01-HD116735 (JS), R01-HD112474 (MCS and RJT) and R01-HD107034 (MCS), by the Department of Defense grant number HT94252310327 (MCS and RJT) and by the NIH National Center for Advancing Translational Sciences through grant number UL1TR001998 (JS, NS). The content is solely the responsibility of the authors and does not necessarily represent the official views of the NIH.

## References

1. Saul, D. et al. Bone Healing Gone Wrong: Pathological Fracture Healing and Non-Unions—Overview of Basic and Clinical Aspects and Systematic Review of Risk Factors. Bioengineering 10, (2023).

2. Wildemann, B. et al. Non-union bone fractures. Nat. Rev. Dis. Primer 7, 1–21 (2021).

3. Kocic, M., Lazovic, M., Mitkovic, M. & Djokic, B. Clinical significance of the heterotopic ossification after total hip arthroplasty. Orthopedics 33, 16 (2010).

4. Alkindi, M. et al. Guided bone regeneration with osteoconductive grafts and PDGF: A tissue engineering option for segmental bone defect reconstruction. J. Appl. Biomater. Funct. Mater. 19, 2280800020987405 (2021).

5. Mahardawi, B., Tompkins, K. A., Mattheos, N., Arunjaroensuk, S. & Pimkhaokham, A. Periosteum-derived Micrografts for bone regeneration. Connect. Tissue Res. 64, 400–412 (2023).

6. Nicholson, J. A., Makaram, N., Simpson, A. & Keating, J. F. Fracture nonunion in long bones: A literature review of risk factors and surgical management. Injury 52 **Suppl 2**, S3–S11 (2021).

7. Łuczak, J. W. et al. The Future of Bone Repair: Emerging Technologies and Biomaterials in Bone Regeneration. Int. J. Mol. Sci. 25, 12766 (2024).

8. Raggatt, L. J. et al. Fracture healing via periosteal callus formation requires macrophages for both initiation and progression of early endochondral ossification. Am. J. Pathol. 184, 3192–3204 (2014).

9. Alexander, K. A. et al. Osteal macrophages promote in vivo intramembranous bone healing in a mouse tibial injury model. J. Bone Miner. Res. Off. J. Am. Soc. Bone Miner. Res. 26, 1517–1532 (2011).

10. Batoon, L. et al. CD169+ macrophages are critical for osteoblast maintenance and promote intramembranous and endochondral ossification during bone repair. Biomaterials 196, 51–66 (2019).

11. Vi, L. et al. Macrophages promote osteoblastic differentiation in-vivo: implications in fracture repair and bone homeostasis. J. Bone Miner. Res. Off. J. Am. Soc. Bone Miner. Res. 30, 1090–1102 (2015).

12. Simkin, J. et al. Macrophages are required to coordinate mouse digit tip regeneration. Dev. Camb. Engl. 144, 3907–3916 (2017).

13. Godwin, J. W., Pinto, A. R. & Rosenthal, N. A. Macrophages are required for adult salamander limb regeneration. Proc. Natl. Acad. Sci. U. S. A. 110, 9415–9420 (2013).

14. Champagne, C. M., Takebe, J., Offenbacher, S. & Cooper, L. F. Macrophage cell lines produce osteoinductive signals that include bone morphogenetic protein-2. Bone 30, 26–31 (2002).

15. Gong, L., Zhao, Y., Zhang, Y. & Ruan, Z. The Macrophage Polarization Regulates MSC Osteoblast Differentiation in vitro. Ann. Clin. Lab. Sci. 46, 65–71 (2016).

16. Chang, M. K. et al. Osteal tissue macrophages are intercalated throughout human and mouse bone lining tissues and regulate osteoblast function in vitro and in vivo. J. Immunol. 181, 1232–1244 (2008).

17. Honda, Y. et al. Elevated extracellular calcium stimulates secretion of bone morphogenetic protein 2 by a macrophage cell line. Biochem. Biophys. Res. Commun. 345, 1155–1160 (2006).

18. Loi, F. et al. The effects of immunomodulation by macrophage subsets on osteogenesis in vitro. Stem Cell Res. Ther. 7, 15 (2016).

19. Zhang, Y. et al. Macrophage type modulates osteogenic differentiation of adipose tissue MSCs. Cell Tissue Res. 369, 273–286 (2017).

20. Wang, T., Zhang, X. & Bikle, D. D. Osteogenic Differentiation of Periosteal Cells During Fracture Healing. J. Cell. Physiol. 232, 913–921 (2017).

21. 21. Shi, L., et al. Identification of novel macrophages and bone morphogenetic protein signals causing ectopic calcification and impairing muscle regeneration. iScience 28, (2025).

22. Yin, W. et al. Activin A secretion by muscle-repairing macrophages induces heterotopic ossification in mice. J. Clin. Invest. 136, (2026).

23. Convente, M. R. et al. Depletion of Mast Cells and Macrophages Impairs Heterotopic Ossification in an Acvr1R206H Mouse Model of Fibrodysplasia Ossificans Progressiva. J. Bone Miner. Res. Off. J. Am. Soc. Bone Miner. Res. 33, 269–282 (2018).

24. Tu, B. et al. Macrophage-Derived TGF-β and VEGF Promote the Progression of Trauma-Induced Heterotopic Ossification. Inflammation 46, 202–216 (2023).

25. Salazar, V. S., Gamer, L. W. & Rosen, V. BMP signalling in skeletal development, disease and repair. Nat. Rev. Endocrinol. 12, 203–221 (2016).

26. Sampath, T. K. & Reddi, A. H. Discovery of bone morphogenetic proteins - A historical perspective. Bone 140, 115548 (2020).

27. James, A. W. et al. A Review of the Clinical Side Effects of Bone Morphogenetic Protein-2. Tissue Eng. Part B Rev. 22, 284–297 (2016).

28. Dawson, L. A. et al. The periosteal requirement and temporal dynamics of BMP2induced middle phalanx regeneration in the adult mouse. Regeneration 4, 140–150 (2017).

29. Howard, M. T. et al. Sustained release of BMP-2 using self-assembled layer-by-layer film-coated implants enhances bone regeneration over burst release. Biomaterials 288, 121721 (2022).

30. Mumcuoglu, D. et al. Injectable BMP-2 delivery system based on collagen-derived microspheres and alginate induced bone formation in a time- and dose-dependent manner. Eur. Cell. Mater. 35, 242–254 (2018).

31. Betz, O. B. et al. Delayed administration of adenoviral BMP-2 vector improves the formation of bone in osseous defects. Gene Ther. 14, 1039–1044 (2007).

32. Yu, Y. Y. et al. Immunolocalization of BMPs, BMP antagonists, receptors, and effectors during fracture repair. Bone 46, 841–851 (2010).

33. Zheng, Z.-Y. et al. Fatty acids derived from apoptotic chondrocytes fuel macrophages FAO through MSR1 for facilitating BMSCs osteogenic differentiation. Redox Biol. 53, 102326 (2022).

34. Ramachandran, P. et al. Resolving the fibrotic niche of human liver cirrhosis at single cell level. Nature 575, 512–518 (2019).

35. Gibbings, S. L. et al. Three Unique Interstitial Macrophages in the Murine Lung at Steady State. Am. J. Respir. Cell Mol. Biol. 57, 66–76 (2017).

36. Aran, D. et al. Reference-based analysis of lung single-cell sequencing reveals a transitional profibrotic macrophage. Nat. Immunol. 20, 163–172 (2019).

37. Simkin, J. et al. Tissue-resident macrophages specifically express Lactotransferrin and Vegfc during ear pinna regeneration in spiny mice. Dev. Cell 59, 496–516.e6 (2024).

38. Lai, S.-L. et al. Reciprocal analyses in zebrafish and medaka reveal that harnessing the immune response promotes cardiac regeneration. eLife 6, e25605 (2017).

39. Chakarov, S. et al. Two distinct interstitial macrophage populations coexist across tissues in specific subtissular niches. Science 363, eaau0964 (2019).

40. Knipper, J. A. et al. Interleukin-4 Receptor α Signaling in Myeloid Cells Controls Collagen Fibril Assembly in Skin Repair. Immunity 43, 803–816 (2015).

41. Simkin, J., Han, M., Yu, L., Yan, M. & Muneoka, K. The mouse digit tip: from wound healing to regeneration. Methods Mol. Biol. Clifton NJ 1037, 419–435 (2013).

42. Fernando, W. A. et al. Wound healing and blastema formation in regenerating digit tips of adult mice. Dev. Biol. 350, 301–310 (2011).

43. Borgens, R. B. Mice regrow the tips of their foretoes. Science 217, 747–750 (1982).

44. Singer, M., Weckesser, E. C., G\’ eraudie, J., Maier, C. E. & Singer, J. Open finger tip healing and replacement after distal amputation in rhesus monkey with comparison to limb regeneration in lower vertebrates. Anat. Embryol. (Berl.) 177, 29–36 (1987).

45. Illingworth, C. M. Trapped fingers and amputated finger tips in children. J. Pediatr. Surg. 9, 853–858 (1974).

46. Neufeld, D. A. & Zhao, W. Phalangeal regrowth in rodents: postamputational bone regrowth depends upon the level of amputation. Prog. Clin. Biol. Res. **383A**, 243–252 (1993).

47. Neufeld, D. A. & Zhao, W. Bone regrowth after digit tip amputation in mice is equivalent in adults and neonates. Wound Repair Regen. Off. Publ. Wound Heal. Soc. Eur. Tissue Repair Soc. 3, 461–466 (1995).

48. Simkin, J. et al. The mammalian blastema: regeneration at our fingertips. Regen. Oxf. Engl. 2, 93–105 (2015).

49. Chamberlain, C. S. et al. Level-specific amputations and resulting regenerative outcomes in the mouse distal phalanx. Wound Repair Regen. Off. Publ. Wound Heal. Soc. Eur. Tissue Repair Soc. 25, 443–453 (2017).

50. Said, S., Parke, W. & Neufeld, D. A. Vascular supplies differ in regenerating and nonregenerating amputated rodent digits. Anat. Rec. A. Discov. Mol. Cell. Evol. Biol. 278, 443–449 (2004).

51. Storer, M. A. et al. Acquisition of a Unique Mesenchymal Precursor-like Blastema State Underlies Successful Adult Mammalian Digit Tip Regeneration. Dev. Cell 52, 509–524.e9 (2020).

52. Johnson, G. L., Masias, E. J. & Lehoczky, J. A. Cellular Heterogeneity and Lineage Restriction during Mouse Digit Tip Regeneration at Single-Cell Resolution. Dev. Cell 52, 525–540.e5 (2020).

53. Tower, R. J. et al. Spatial transcriptomics reveals metabolic changes underly age-dependent declines in digit regeneration. eLife 11, e71542 (2022).

54. Jin, S. et al. Inference and analysis of cell-cell communication using CellChat. Nat. Commun. 12, 1088 (2021).

55. Browaeys, R., Saelens, W. & Saeys, Y. NicheNet: modeling intercellular communication by linking ligands to target genes. Nat. Methods 17, 159–162 (2020).

56. Jensen, K. B., Driskell, R. R. & Watt, F. M. Assaying proliferation and differentiation capacity of stem cells using disaggregated adult mouse epidermis. Nat. Protoc. 5, 898–911 (2010).

57. Hoang, G. et al. Uncovering metabolic reservoir cycles in MYC-transformed lymphoma B cells using stable isotope resolved metabolomics. Anal. Biochem. 632, 114206 (2021).

58. Trygg, J., Holmes, E. & Lundstedt, T. Chemometrics in metabonomics. J. Proteome Res. 6, 469–479 (2007).

59. Trygg, J. & Wold, S. Orthogonal projections to latent structures (O-PLS). J. Chemom. 16, 119–128 (2002).

60. Wold, S., Sjöström, M. & Eriksson, L. PLS-regression: a basic tool of chemometrics. Chemom. Intell. Lab. Syst. 58, 109–130 (2001).

61. Dang, D. et al. Computational Approach to Identifying Universal Macrophage Biomarkers. Front. Physiol. 11, 275 (2020).

62. Wang, S. et al. Selenoprotein K protects skeletal muscle from damage and is required for satellite cells-mediated myogenic differentiation. Redox Biol. 50, 102255 (2022).

63. Zhang, W. et al. Roles of selenoprotein K in oxidative stress and endoplasmic reticulum stress under selenium deficiency in chicken liver. Comp. Biochem. Physiol. Part C Toxicol. Pharmacol. 264, 109504 (2023).

64. Huang, Z., Rose, A. H. & Hoffmann, P. R. The Role of Selenium in Inflammation and Immunity: From Molecular Mechanisms to Therapeutic Opportunities. Antioxid. Redox Signal. 16, 705–743 (2012).

65. Misra, S. et al. Loss of selenoprotein W in murine macrophages alters the hierarchy of selenoprotein expression, redox tone, and mitochondrial functions during inflammation. Redox Biol. 59, 102571 (2022).

66. Mills, E. L. et al. Succinate Dehydrogenase Supports Metabolic Repurposing of Mitochondria to Drive Inflammatory Macrophages. Cell 167, 457–470.e13 (2016).

67. Tannahill, G. M. et al. Succinate is an inflammatory signal that induces IL-1β through HIF-1α. Nature 496, 238–242 (2013).

68. Wu, K. K.-L. et al. MDM2 induces pro-inflammatory and glycolytic responses in M1 macrophages by integrating iNOS-nitric oxide and HIF-1α pathways in mice. Nat. Commun. 15, 8624 (2024).

69. Freemerman, A. J. et al. Metabolic reprogramming of macrophages: glucose transporter 1 (GLUT1)-mediated glucose metabolism drives a proinflammatory phenotype. J. Biol. Chem. 289, 7884–7896 (2014).

70. Rodríguez-Prados, J.-C. et al. Substrate fate in activated macrophages: a comparison between innate, classic, and alternative activation. J. Immunol. Baltim. Md 1950 185, 605–614 (2010).

71. Newsholme, P., Gordon, S. & Newsholme, E. A. Rates of utilization and fates of glucose, glutamine, pyruvate, fatty acids and ketone bodies by mouse macrophages. Biochem. J. 242, 631–636 (1987).

72. Newsholme, P., Curi, R., Gordon, S. & Newsholme, E. A. Metabolism of glucose, glutamine, long-chain fatty acids and ketone bodies by murine macrophages. Biochem. J. 239, 121–125 (1986).

73. Seim, G. L. et al. Two-stage metabolic remodelling in macrophages in response to lipopolysaccharide and interferon-γ stimulation. Nat. Metab. 1, 731–742 (2019).

74. Bae, S. et al. MYC-mediated early glycolysis negatively regulates proinflammatory responses by controlling IRF4 in inflammatory macrophages. Cell Rep. 35, 109264 (2021).

75. Lauterbach, M. A. et al. Toll-like Receptor Signaling Rewires Macrophage Metabolism and Promotes Histone Acetylation via ATP-Citrate Lyase. Immunity 51, 997–1011.e7 (2019).

76. Vats, D. et al. Oxidative metabolism and PGC-1beta attenuate macrophage-mediated inflammation. Cell Metab. 4, 13–24 (2006).

77. Huang, S. C.-C. et al. Cell-intrinsic lysosomal lipolysis is essential for macrophage alternative activation. Nat. Immunol. 15, 846–855 (2014).

78. Mills, E. L. & O’Neill, L. A. Reprogramming mitochondrial metabolism in macrophages as an anti-inflammatory signal. Eur. J. Immunol. 46, 13–21 (2016).

79. Zhang, S. et al. Efferocytosis Fuels Requirements of Fatty Acid Oxidation and the Electron Transport Chain to Polarize Macrophages for Tissue Repair. Cell Metab. 29, 443–456.e5 (2019).

80. Yu, L., Han, M., Yan, M., Lee, J. & Muneoka, K. BMP2 induces segment-specific skeletal regeneration from digit and limb amputations by establishing a new endochondral ossification center. Dev. Biol. 372, 263–273 (2012).

81. Lee, J. et al. SDF-1α/CXCR4 signaling mediates digit tip regeneration promoted by BMP-2. Dev. Biol. 382, 98–109 (2013).

82. Yu, L. et al. BMP signaling induces digit regeneration in neonatal mice. Dev. Camb. Engl. 137, 551–559 (2010).

83. Yu, L. et al. BMP9 stimulates joint regeneration at digit amputation wounds in mice. Nat. Commun. 10, 424 (2019).

